# Antiparasitic effect of stilbene and terphenyl compounds against *Trypanosoma cruzi* parasites

**DOI:** 10.1101/2021.02.23.432446

**Authors:** Federica Bruno, Germano Castelli, Fabrizio Vitale, Simone Catanzaro, Valeria Vitale Badaco, Marinella Roberti, Claudia Colomba, Antonio Cascio, Manlio Tolomeo

**Affiliations:** National Reference Center for Leishmaniasis (C.Re.Na.L.), Istituto Zooprofilattico Sperimentale della Sicilia, Via Gino Marinuzzi 3, 90129, Palermo, Italy; Department of Pharmacy and Biotechnology, University of Bologna, Via Belmeloro 6, 40126, Bologna, Italy; Department of Health Promotion Sciences, Section of Infectious Diseases, University of Palermo, Via del Vespro 129, 90127, Palermo, Italy

## Abstract

**Background:** Chagas disease, also known as American trypanosomiasis, is a potentially life-threatening illness caused by the protozoan parasite *Trypanosoma cruzi*. No progress in the treatment of this pathology has been made since Nifurtimox was introduced more than fifty years ago and is considered very aggressive and may cause several adverse effects. Currently, this drug has severe limitations, including high frequency of undesirable side effects and limited efficacy and availability and the research to discover new drugs for the treatment of Chagas disease is imperative. Many drugs available in the market are natural products as found in nature or compounds designed based on the structure and activity of these natural products.

**Methodology/Principal Findings:** This study evaluated the in vitro antiparasitic activity in *T. cruzi* epimastigotes and intracellular amastigotes of a series of stilbene and terphenyl compounds previously synthesized. The action of the most selective compounds has been investigated by flow cytometry analysis to evaluate the mechanism of cell death. The ability to induce apoptosis or caspase-1 inflammasome were assayed in macrophages infected with *T. cruzi* after treatment comparing with Nifurtimox.

**Conclusions/Significance:** The stilbene ST18 was the most potent compound of the series. It was slightly less active than Nifurtimox in epimastigotes but most active in intracellular amastigotes. Compared to Nifurtimox, it was markedly less cytotoxic when tested in vitro on normal cells. ST18 was able to induce a marked increase of parasites positive to Annexin V and monodansylcadaverine. Moreover, ST18 induced the activation in infected macrophages of caspase-1, a conserved enzyme which plays a main role in controlling parasitemia, host survival, and the onset of adaptive immune response in Trypanosoma infection. The antiparasitic activity of ST18 together to its ability to activate caspase-1 in infected macrophages and its low toxicity on normal cells makes this compound interesting for further clinical investigations.

**Author Summary:** Chagas disease is a pathology caused by the protozoan parasite *Trypanosoma cruzi.* No progress in the treatment of this pathology has been made since benznidazole and Nifurtimox were introduced more than fifty years ago. However, these drugs have severe limitations and the research to discover new drugs for the treatment of Chagas disease is imperative. We evaluated the *in vitro* antiparasitic activity in *T. cruzi* epimastigotes of a series of stilbene and terphenyl compounds previously synthesized. The stilbene ST18 was the most potent compound of the series. It was slightly less active than nifurtimox in epimastigotes but most active in intracellular amastigotes. Compared to Nifurtimox, it was markedly less cytotoxic when tested in vitro on normal cells. ST18 was able to induce a marked increase of parasites positive to Annexin V and monodansylcadaverine. Moreover, this compound induced the activation in infected macrophages of caspase-1, an evolutionarily conserved enzyme which plays a main role in controlling parasitemia, host survival, and the onset of adaptive immune response in *T. cruzi* infection. The antiparasitic activity of ST18 together to its ability to activate caspase-1 in infected macrophages and its low toxicity on normal cells makes this compound interesting for further clinical investigations.

## Introduction

*Trypanosoma cruzi* (*T. cruzi*) is a protozoan parasite transmitted primarily by triatomine insects. It is the agent of Chagas disease, an endemic pathology in Latin America that affects about 6-8 million people worldwide [1] and causes approximately 50,000 deaths per year. Only two nitroheterocyclic drugs, Nifurtimox and benznidazole, are available for the treatment of Chagas disease. Currently, these drugs have severe limitations, including high frequency of undesirable side effects, long protocols of treatment, and limited efficacy and availability; although they are effective for the treatment of acute infections. Experimental toxicity studies with Nifurtimox evidenced neurotoxicity, testicular damage, ovarian toxicity, and deleterious effects in adrenal, colon, oesophageal and mammary tissue which frequently necessitate the cessation of treatment. In the case of benznidazole deleterious effects were observed in adrenals, colon and oesophagus. Both drugs exhibited significant mutagenic effects and were shown to be tumorigenic or carcinogenic in some studies [2,3]. Therefore, natural products have always been a source of a great variety of bioactive molecules, mostly substances from the organism secondary metabolism. Many drugs available in the market are natural products as found in nature or compounds designed based on the structure and activity of these natural products (semi-synthetic or completely synthetic)[4]. Recently, several natural and synthetic stilbene and terphenyl have been studied for their anticancer and leishmanicidal properties [5–8], in particular we evaluated the antileishmanial activity of two compounds, a trans-stilbene derivatives and a terphenyl derivatives, namely trans-1,3-dimethoxy-5-(4-methoxystyryl) benzene (ST18) and 3,4”,5-trimethoxy-1,1’:2’,1”-terphenyl (TR4), presented the best activity and safety profiles [9,10].

In the current study we evaluated the *in vitro* antiparasitic activity in *T. cruzi* epimastigotes of a series of *cis*- and *trans*-stilbene derivatives in which a variety of substituents were introduced at position 2’, 3’ and 4’ of the stilbene scaffold while the 3-5dimethoxy motif was maintained. Additionally, we studied a series of terphenyl compounds incorporating a phenyl ring as a bioisosteric substitution of the stilbene alkenyl bridge.

We observed that the stilbene ST18 was endowed with potent antiparasitic activity in both *T. cruzi* epimastigotes and intracellular *T. cruzi* amastigotes. Compared to Nifurtimox, it was markedly less cytotoxic when tested *in vitro* on normal and differentiated cells. Moreover, this compound induced the activation in infected macrophages of caspase-1, an evolutionarily conserved enzyme which plays a main role for controlling parasitemia, host survival, and the onset of adaptive immune response in *T. cruz*i infection.

## Materials and Methods

### Parasites cultures

A strain of *T. cruzi* taken by stock archive of the OIE Reference Laboratory National Reference Center for Leishmaniasis (C.Re.Na.L. Palermo, Italy) was cultured in 25 cm^2^ flasks (Falcon) at 25 °C and pH 7.18 in RPMI-PY medium, which consisted of RPMI 1640 (Sigma R0883) supplemented with equal volume of Pepton-yeast medium, 10% fetal bovine serum (FBS), 1% glutamine, 250 μg/mL gentamicin and 500 μg/mL of 5-fluorocytosine [11].

### Compounds and sample preparation

Compounds ST18 and 6 were synthesized as reported by Kim et al. [12]; compounds 1-5 and 8-10 were prepared as previously described by us [6], compounds 7, TR4 and 13-14 were prepared as previously described by us [7], 15 was synthesized as reported by Pizzirani et al. [5]. The purity of compound was determined by elemental analyses and was ≥ 97%. Each compound was dissolved in dimethyl sulfoxide (DMSO) in a stock solution at a concentration of 20 mM, stored at −20°C and protected from light. In each experiment DMSO never exceeded 0.2% and this percentage did not interfere with cell growth. Nifurtimox was obtained from Merck Sigma-Aldrich (Milano, Italy).

### Epimastigotes viability assay

To evaluate the effects of compounds in cultures of *T. cruzi* a viability assay protocol similar to that described by Castelli et al. [9] was used with some modifications. Exponentially growing of *T. cruzi* were dispensed at the concentration of 4×10^6^/mL in 25-m2 flasks (Falcon) and treated with increasing concentrations (from 1 to 200 μM) of each compound. After 72 h of treatment the parasites were centrifugated and resuspended in 1 ml of RPMI-PY medium. The suspension of *T. cruzi* from each treatment was mixed with 0.4% trypan blue solution at a ratio of 3:1 (vol/vol). The percentage of vitality of *T. cruzi* was observed by counting in a Bürker hemocytometer for enumeration of stained and unstained cells, taken respectively as dead and living cells, in comparison with the control culture (100% viability). IC_50_ (half maximal inhibitory concentration) was evaluated after 72 h and was calculated by regression analysis (GraphPad software).

### Effects of compounds in intracellular amastigotes

U937 monocytic cells (1×10^5^ cells/mL) in the logarithmic phase of growth were plated onto chamber Lab Tek culture slides in 2.5 mL of RPMI 1640 (Sigma), 10% FBS medium containing 25 ng/mL of phorbol 12-myristate 13-acetate (Sigma) for 18 h to induce macrophage differentiation.

After incubation, the medium was removed by washing twice with RPMI-1640 medium. Non adherent cells were removed, and the macrophages were further incubated overnight in RPMI 1640 medium supplemented with 10% FBS. Then adherent macrophages were infected with *T. cruzi* epimastigotes at a parasite/macrophage ratio of 50:1 for 24 h at 37 °C in 5% CO_2_. Free epimastigotes were removed by three extensive washing with RPMI 1640 medium, and infected macrophages were either incubated 48 h in media alone (control) or with Nifurtimox, ST18 or TR4. With the aim of stain intracellular amastigotes, cells were fixed with iced methanol to permeabilize cell membrane to ethidium bromide and stained with 100 μg/mL ethidium bromide. The number of amastigotes was determined by examining three coverslips for each treatment. At least 200 macrophages were counted by visual examination under 400× magnifications by using a fluorescence microscope Nikon Eclipse E200 (Nikon Instruments Europe, Amsterdam, Netherlands) equipped with a green filter to determine the number of intracellular amastigotes. The number of intracellular amastigotes in samples treated with each compound was expressed as percentage of the untreated control.

### Mammalian cell cytotoxicity

Potential cytotoxic action of each compound was checked by 3-(4,5-dimethylthiazol-2-yl)-2,5-diphenylterazolium bromide (MTT) assay on macrophages derived by U937 cells and in primary epithelial cells of Cercopiteco (CPE). Macrophages and CPE cells were cultured in RPMI 1640 (Sigma) supplemented with 10% fetal bovine serum (FBS, Gibco), penicillin (100 IU/mL) and streptomycin (100 mg/mL). Cells were grown at 37 °C in 5% CO_2_ and passaged twice a week. In each experiment, cells (10^5^/well) were incubated into 96-well plates overnight in a humidified 5% CO_2_ atmosphere at 37 °C to ensure cell adherence. After 24 h, cells were treated with increasing concentrations of each compound. Non-treated cells were included as a negative control. After 72 h incubation with each compound, the MTT (5 mg/mL) was added to each well and incubated at 37 °C for 4 h. Then the medium and MTT were removed, cells washed by PBS and 200 μL of DMSO were added to dissolve the formazan crystals. The absorbance was measured using a microplate reader Spectrostar Nano (BMG LabTech) at 570 nm. The reduction of MTT to insoluble formazan was done by the mitochondrial enzymes of viable cells and so is an indicator of cell viability. Therefore, decreases of absorbance indicate toxicity to the cell. The viability was calculated using the following formula: [(L2/L1) × 100], where L1 is the absorbance of control cells and L2 is the absorbance of treated cells. The IC50 was calculated by regression analysis (GraphPad software).

The selectivity index (SI) was determined by dividing the IC50 calculated in mammalian cells and the IC50 calculated in *T. cruzi* parasites.

### Cell cycle analysis by flow cytometry

Epimastigotes (4×10^6^) were incubated for 48 h with each compounds at 26 °C. Afterward, parasites were washed 3 times with PBS containing 0.02 M EDTA to avoid clumps and were then fixed with cold methanol for 24 h. The parasites were resuspended in 0.5 mL of PBS containing RNase I (50 μg/mL) and PI (25 μg/mL) and were then incubated at 25 °C for 20 min. The material was kept on ice until analysis. The stained parasites were analysed in single-parameter frequency histograms by using a FACScan flow cytometer (Becton Dickinson, CA).

### Cell volume determination

Epimastigotes were collected by centrifugation at 1,000g, washed twice in PBS, resuspended in PBS to 500×10^3^ parasites/mL, and analysed by FACScan flow cytometer (Becton Dickinson, CA). Density plots of forward (FSC) versus side (SSC) scatter represent the acquisition of 10×10^3^ events.

### Determination of apoptosis by Annexin V

Externalization of phosphatidylserine on the outer membrane of parasites with and without treatment was determined by using Annexin V labeling kit following the manufacturer’s protocol (Annexin-V-FITC Apoptosis Detection Kit Alexis, Switzerland). Briefly, epimastigotes (2×10^6^) were washed with PBS and centrifuged at 500 g for 5 min. The pellet was suspended in 100 μL of staining solution containing FITC-conjugated Annex-in-V and propidium iodide (Annexin-V-Fluos Staining Kit, Roche Molecular Biochemicals, Germany) and incubated for 15 min at 20 °C. Annexin V positive parasites were determined by using a FACScan flow cytometer (Becton Dickinson, CA).

### Monodansylcadaverine labelling

Monodansylcadaverine (MDC), which is an autofluorescent compound due to the dansyl residue conjugated to cadaverine, have been shown to accumulate in acidic autophagic vacuoles. The concentration of MDC in autophagic vacuole is the consequence of an ion-trapping mechanism and an interaction with lipids in autophagic vacuoles (autophagic vacuoles are rich in membrane lipids).

The use of MDC staining is a rapid and convenient approach to assay autophagy, as shown in cultured cells [13]. Autophagic vacuoles were labeled with MDC by incubating cells on coverslips with 0.05 mM MDC in PBS at 37°C for 10 minutes. After incubation, cells were washed four times with PBS and immediately analysed by fluorescence microscopy (Nikon Eclipse E 200, Japan) equipped with a blue filter. Images were obtained with a Nikon Digital Sight DS-SM (Nikon, Japan) camera and processed using the program EclipseNet, version 1.20.0 (Nikon, Japan).

### Caspase-1 detection

To evaluate the level of active caspase-1, U937 cell line in macrophagic form infected with *T. cruzi* was used. Infected macrophages were incubated for 24 h at 37 °C in 5% CO_2_. Free parasites were removed by extensive washing with RPMI-1640 medium, and infected cells were either incubated in media alone (infection control) or with each compound. After 48h, the culture medium was removed and treated with caspase-1 assay kit (Promega) following the manufacturer’s instructions.

### Statistical analysis

All assays were performed by two observers in three replicates samples and repeated with three new batches of parasites. The mean and standard error of at least three experiments were determined. The differences between the mean values obtained for experimental groups were evaluated by the Student’s t test. P-values of 0.05 or less were considered significant. All statistical analysis was performed using GraphPad Prism 5 software. The IC50 values were calculated by linear regression.

## Results

### Anti-*Trypanosoma cruzi* activity

Table 1 shows the *in vitro* antiparasitic effects evaluated as IC_50_ of different stilbenes (ST18, 1-10) and terphenyls (TR4, 11-15) in *T. cruzi* epimastigotes. These compounds were previously synthesized by us except ST18 and 6 that were reported by Kim et al. [12]. Data were compared to those obtained with Nifurtimox which is the drug currently used for the treatment of *T. cruzi* infection. The most active compounds of the series were the stilbene ST18 (IC50 = 4.6 μM) and the terphenyl TR4 (IC50 = 30 μM).

Figure 1A shows the *in vitro* effects of Nifurtimox, ST18 and TR4 used at increasing concentrations for 72 h in *T. cruzi* epimastigotes. ST18 was markedly more potent than TR4 but less active than Nifurtimox. Upon entering the mammalian host, *T. cruzi* parasites transform into the amastigote stage that reside inside the phagolysosomal vacuoles of macrophages. We evaluated the anti-amastigote efficacy in differentiated macrophage cells (derived from U937 cells) infected with *T. cruzi* as reported in material and methods. Infected macrophages were treated with Nifurtimox, ST18 and TR4 used at increasing concentrations for 72 h. Differently from the results obtained in epimastigotes, the anti-parasitic effect of ST18 in infected macrophages was higher than that observed using Nifurtimox (Fig. 1B).

**Fig 1.**
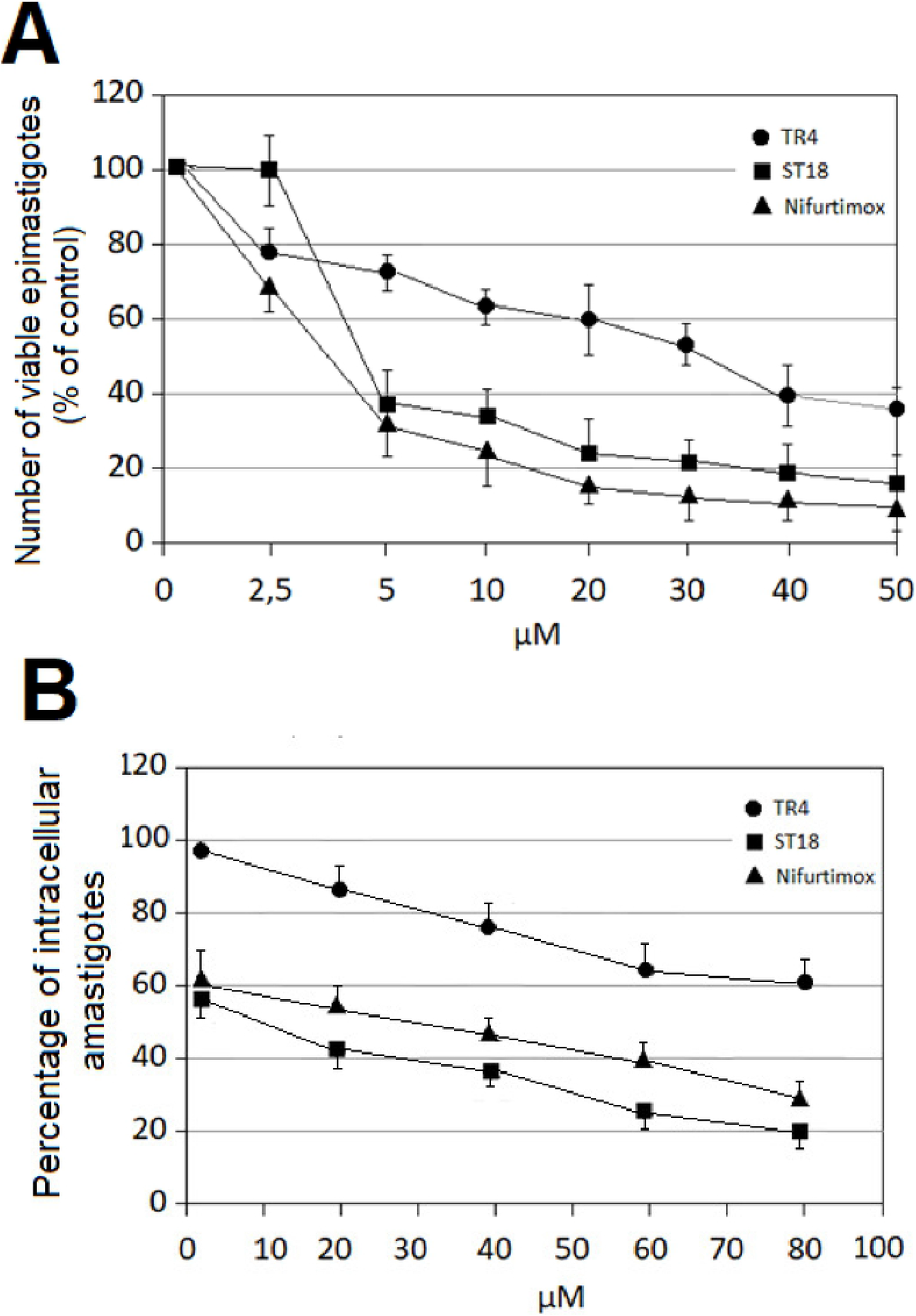
Effects of compounds Nifurtimox, ST18 and TR4 in *Trypanosoma cruzi* epimastigotes and intracellular amastigotes. **(A)** Number of viable *T. cruzi* epimastigotes expressed as percentage of untreated control after 72 h exposure to increasing concentrations of Nifurtimox, ST18 and TR4. **(B)** Number of intracellular amastigotes expressed as percentage of the untreated control after 72 h treatment with Nifurtimox, ST18 and TR4. Bars indicate the mean ± SE from four independent experiments. Data obtained are statistically significant at P < 0.05.

### Mammalian cell cytotoxicity and SI

Primary epithelial cells of Cercopiteco (CPE) and macrophages derived by differentiation of U937 cells were treated with increasing concentrations of ST18 and Nifurtimox. Cytotoxicity was evaluated after 72 h through MTT assay. ST18 showed a very low cytotoxicity in both cell lines compared to Nifurtimox (Fig. 2A) In macrophages the IC50 of ST18 was 143 μM while the IC50 of Nifurtimox was 28 μM with a SI of 31 for ST18 and 8.75 for Nifurtimox. In CPE the IC50s of ST18 and Nifurtimox were 155 μM and 77 μM respectively with a SI of 33.7 for ST18 and 24 for Nifurtimox. (Fig. 2B)

**Fig 2.**
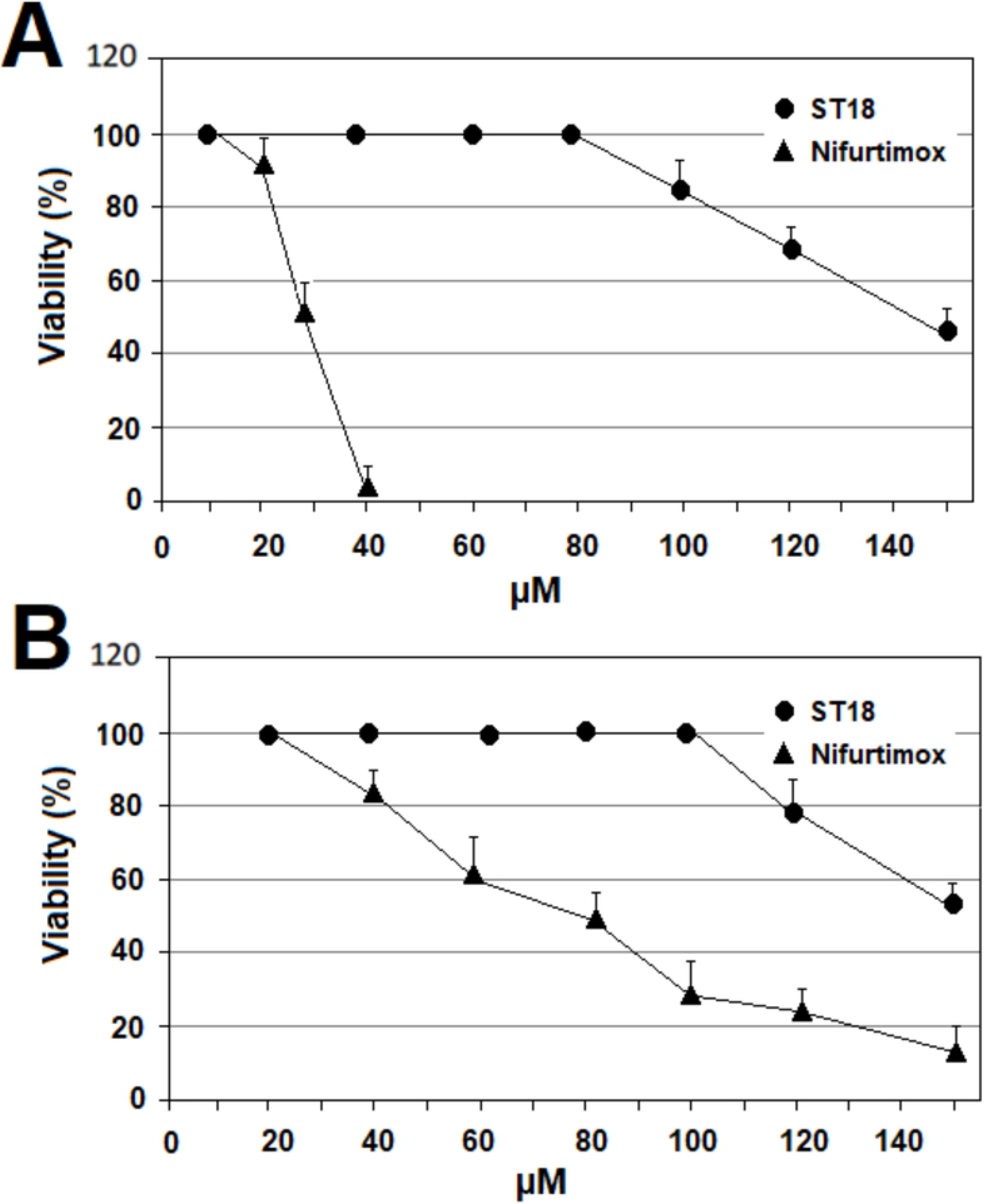
Cytotoxic effects of compounds ST18 and Nifurtimox in Mammalian cells. **(A)** Cytotoxic effects of compounds ST18 and Nifurtimox in primary epithelial cells of Cercopiteco (CPE). **(B)** Cytotoxic effects of compounds ST18 and Nifurtimox in U937 macrophage cells. Bars indicate the mean ± SE from four independent experiments. Data obtained are statistically significant at P< 0.05.

### Cell cycle

The effects of Nifurtimox, ST18 and TR4 on cell cycle distribution of *T. cruzi* was analysed by FACScan flow cytometer. To exclude in the study of cell cycle dead cells that are often located in a sub-G0-G1 peak we decided to study the effects of each compound on the cell cycle by treating the parasites for a period of time and with concentrations of each compound that caused a block of cell growth (evaluated by counting parasites on a hemocytometer) without causing a relevant number of dead cells (evaluated by trypan blue staining). Since after 72 h of treatment the cell growth inhibition was associated to an increase in cell death number (data not shown) we studied the effects of each compound on cell cycle after only 48 h of drug exposure treating parasites with 35 μM Nifurtimox, 50 μM ST18 and 90 μM TR4. This treatment caused a complete block of cell growth with a percentage of dead cells lower than 10%. Cell cycle distribution was analysed by the standard propidium iodide procedure. Nifurtimox did not determined important variations in cell cycle distribution, but only a little reduction in the G2M peack. In contrast, ST18 caused an evident block in G2M while TR4 a block in G1 (Fig 3).

**Fig 3.**
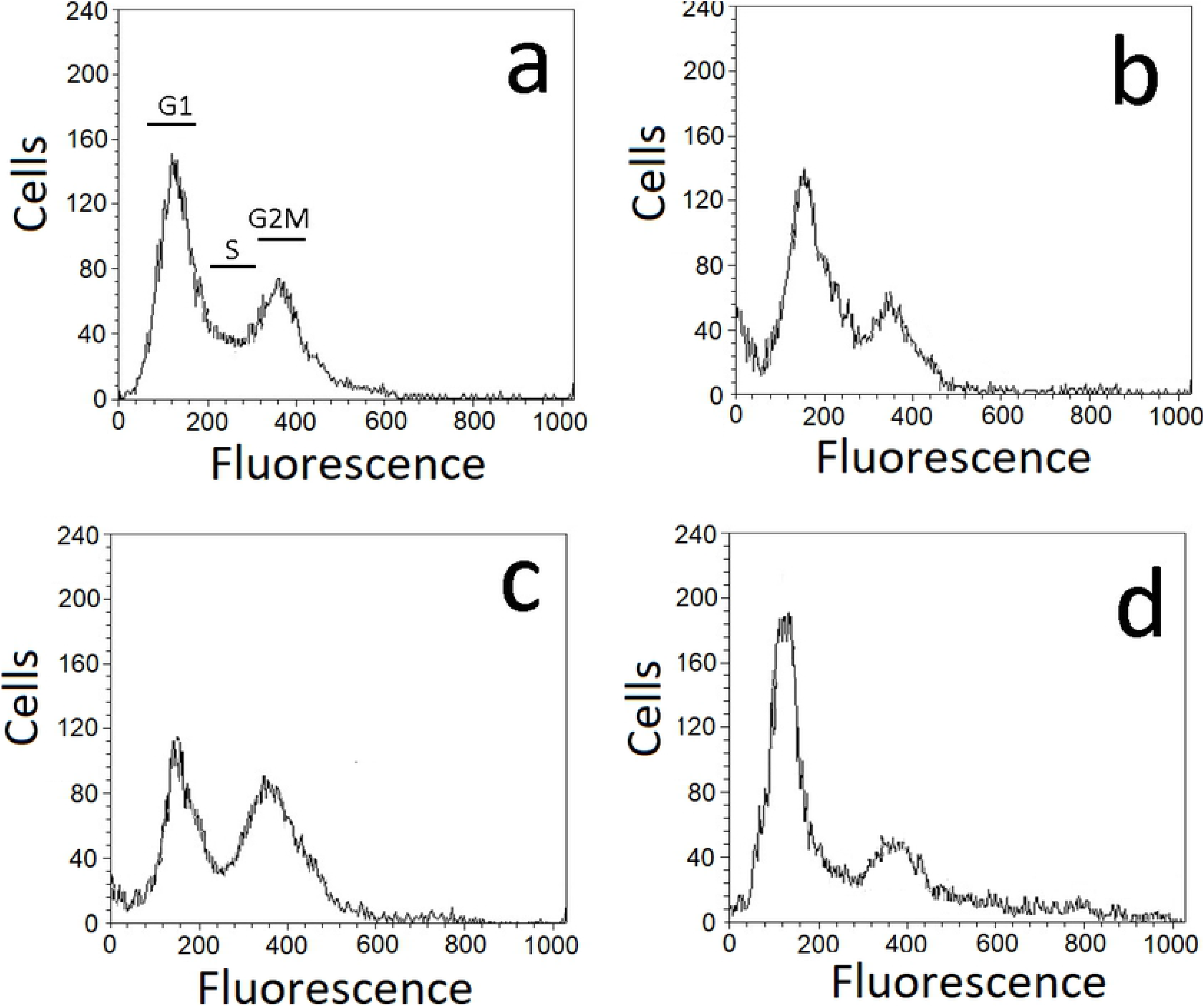
Effects of Nifurtimox, ST18 and TR4 on DNA content/parasite in *Trypanosoma cruzi* epimastigotes. The parasites were cultured without compound (control, panel a) or with 35 μM of Nifurtimox (panel b), 50 μM of ST18 (panel c) and 90 μM of TR4 (panel d). Cell cycle distribution was analysed by the standard propidium iodide procedure. G1, S, and G2–M cells are indicated in panel a.

### Physical parameters

We studied the physical parameters of *T. cruzi* parasites treated with Nifurtimox, ST18 and TR4 by FACScan flow cytometer as previously reported by Jimenez et al [14]. Figure 4A shows density plots for forward scatter (FSC) versus side scatter (SSC) in *T. cruzi* epimastigotes untreated or treated with with 35 μM Nifurtimox, 50 μM ST18, and 90 μM TR4 for 72 h. The measurement of forward scatter allows for the discrimination of cells by size. FSC intensity is proportional to the diameter of the cell. Side scatter measurement provides information about the internal complexity (i.e. granularity) of a cell. The analysis of the density plot of Trypanosome epimastigotes treated with Nifurtimox shows a marked reduction in the average cell size compared to the control. In contrast, FACS analysis of Trypanosome epimastigotes treated with ST18 shows a heterogeneous population characterized by parasites with low dimension and parasites with increased size and granularity. No important modifications were observed with TR4. These data were confirmed by the FACS histograms as shown in Figure 4B.

**Fig 4.**
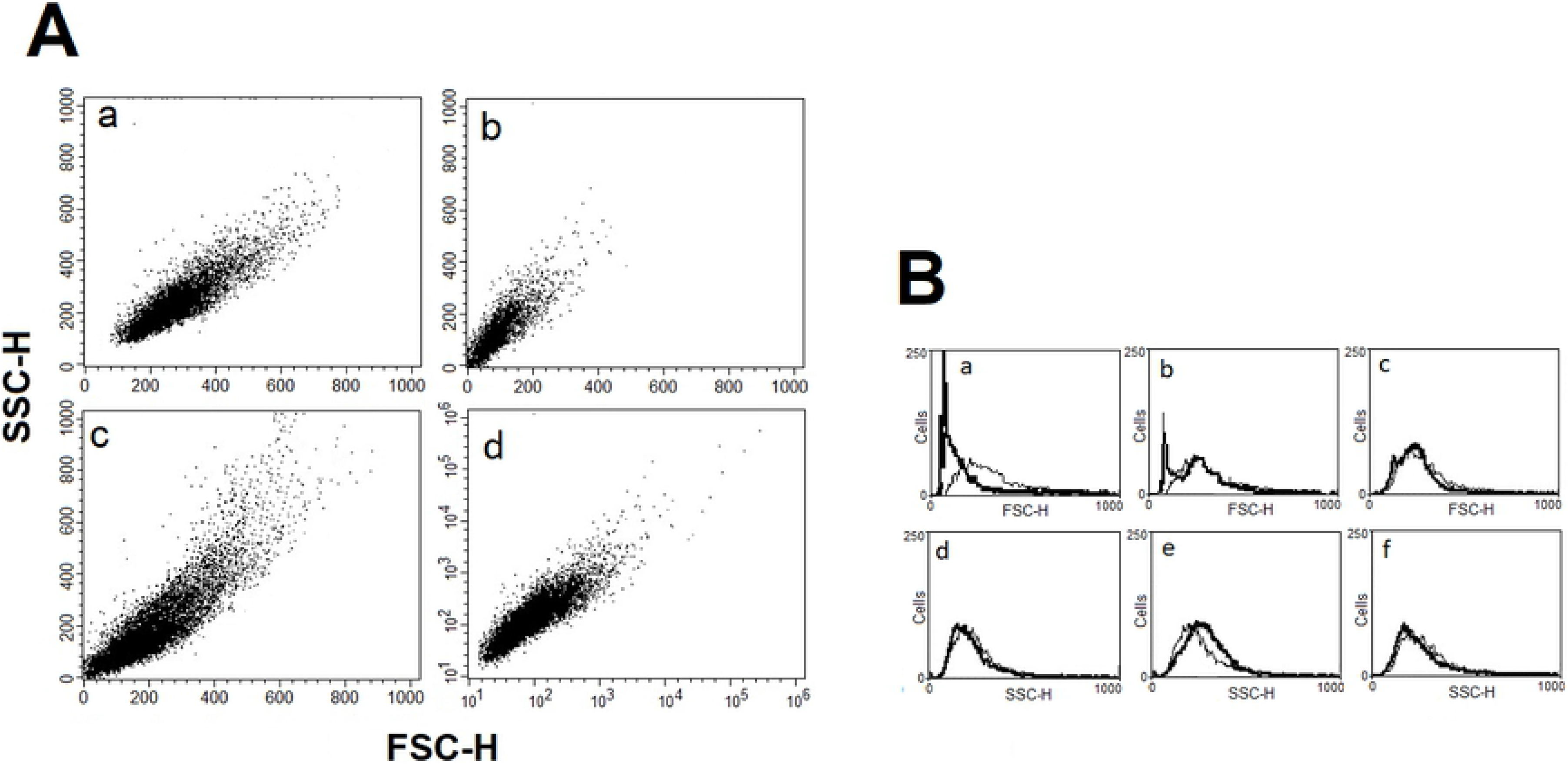
FACS analysis of *Trypanosoma cruzi* epimastigotes cell volume populations. **(A)** Forward light scatter (FSC-H) was considered as function of cell size and side light scatter (SSC-H) as result of cell granularity. Density plots for FSC versus SSC in *T. cruzi* epimastigotes after 72 h treatment with Nifurtimox (panel b), ST18 (panel c) and TR4 (panel d). Untreated control is represented in panel a. **(B)** Representative FACS histogram showing FSC-H and SSC-H of *T. cruzi* epimastigotes after 72 h treatment with Nifurtimox (panels a and d) ST18 (panels b and e) and TR4 (panels c and f). Thin line: No treated control parasites; thick line: parasites treated with each compound. Data are representative of three separate experiments.

### Annexin V and MDC labeling

The loss of cell volume or cell shrinkage is a hallmark of the early phase of the apoptotic process. In order to confirm whether volume reduction of parasites was related to apoptosis, the exposition of phosphatidylserine at the cell surface was analysed by Annexin V labeling test after treatment with Nifurtimox, ST18 and TR4. A significant increase in the percentage of parasites positive to Annexin V was observed after treatment with Nifurtimox and, to a lesser extent, after treatment with ST18 (Fig. 5).

**Fig 5.**
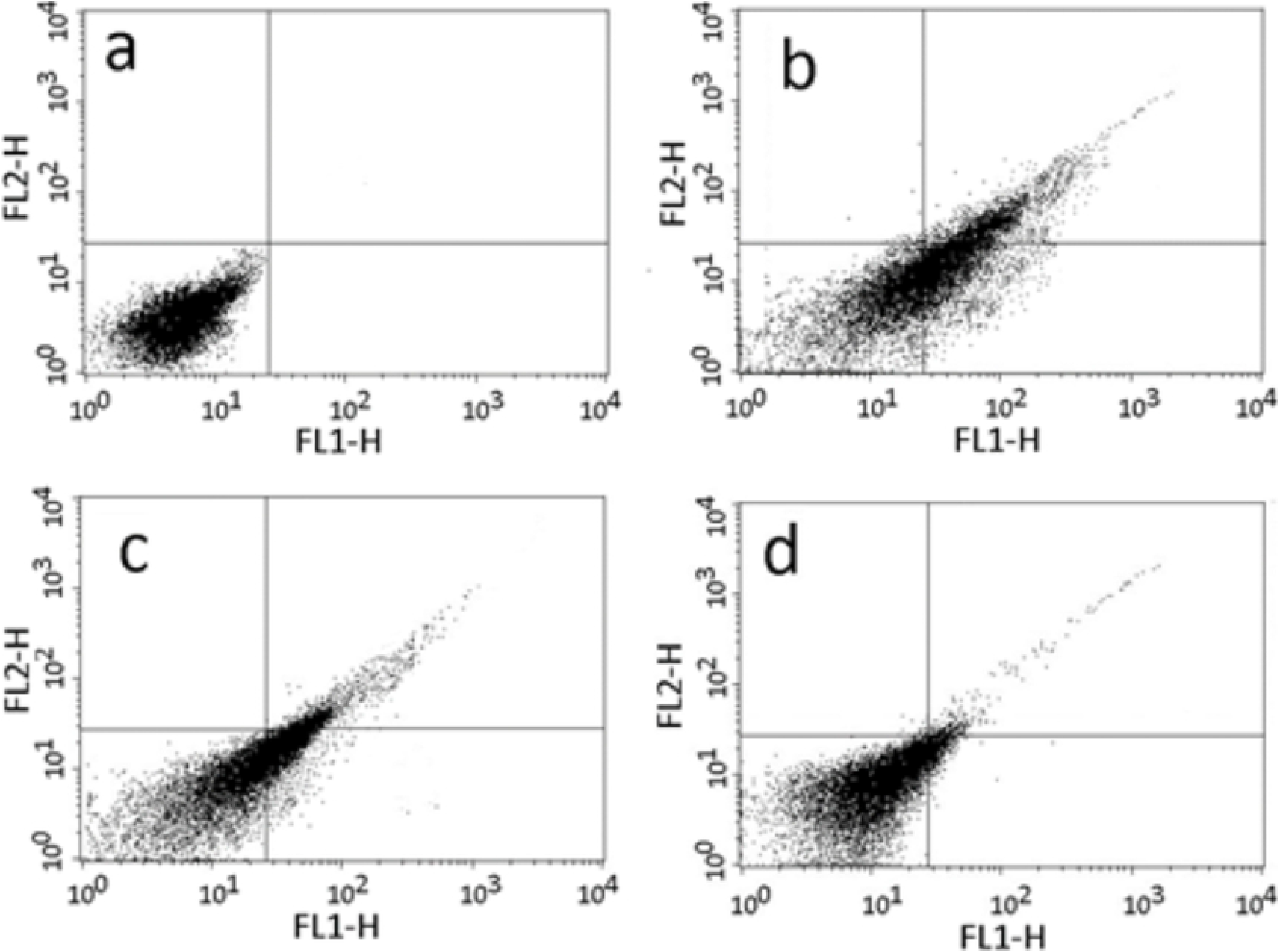
Analysis of phosphatidylserine (PS) extracellular exposure. Representative dot plot of FACS analysis for PS exposure, measured by double staining with Annexin V-FITC and propidium iodide (PI) in *T. cruzi* epimastigotes after 72 h treatment with Nifurtimox (panel b) and ST18 (panel c) and TR4 (panel d). No treated control is represented in panel a. Lower left quadrant belongs to control cells (Annexin V negative/PI negative), lower right quadrant belongs to early apoptotic cells (Annexin V positive/PI negative), upper right quadrant belongs to late apoptotic cells (Annexin V positive/PI positive), upper left quadrant belongs to necrotic cells (Annexin V negative/PI positive). Data are representative of three separate experiments.

Since the analysis of physical parameters of *T. cruzi* treated with ST18 showed also a cell population with increased size and granularity, parameters that are hallmarks of the autophagic process, parasites were treated with monodansylcadaverine (MDC), a specific fluorescent marker for autophagic vacuoles [15]. About 30% of parasites treated 72 h with 40 μM ST18 were strongly positive to MDC test showing numerous fluorescent vacuoles in the cytoplasm. These vacuoles were not observed in untreated control and in samples treated with Nifurtimox or TR4 (data not shown) (Fig. 6).

**Fig 6.**
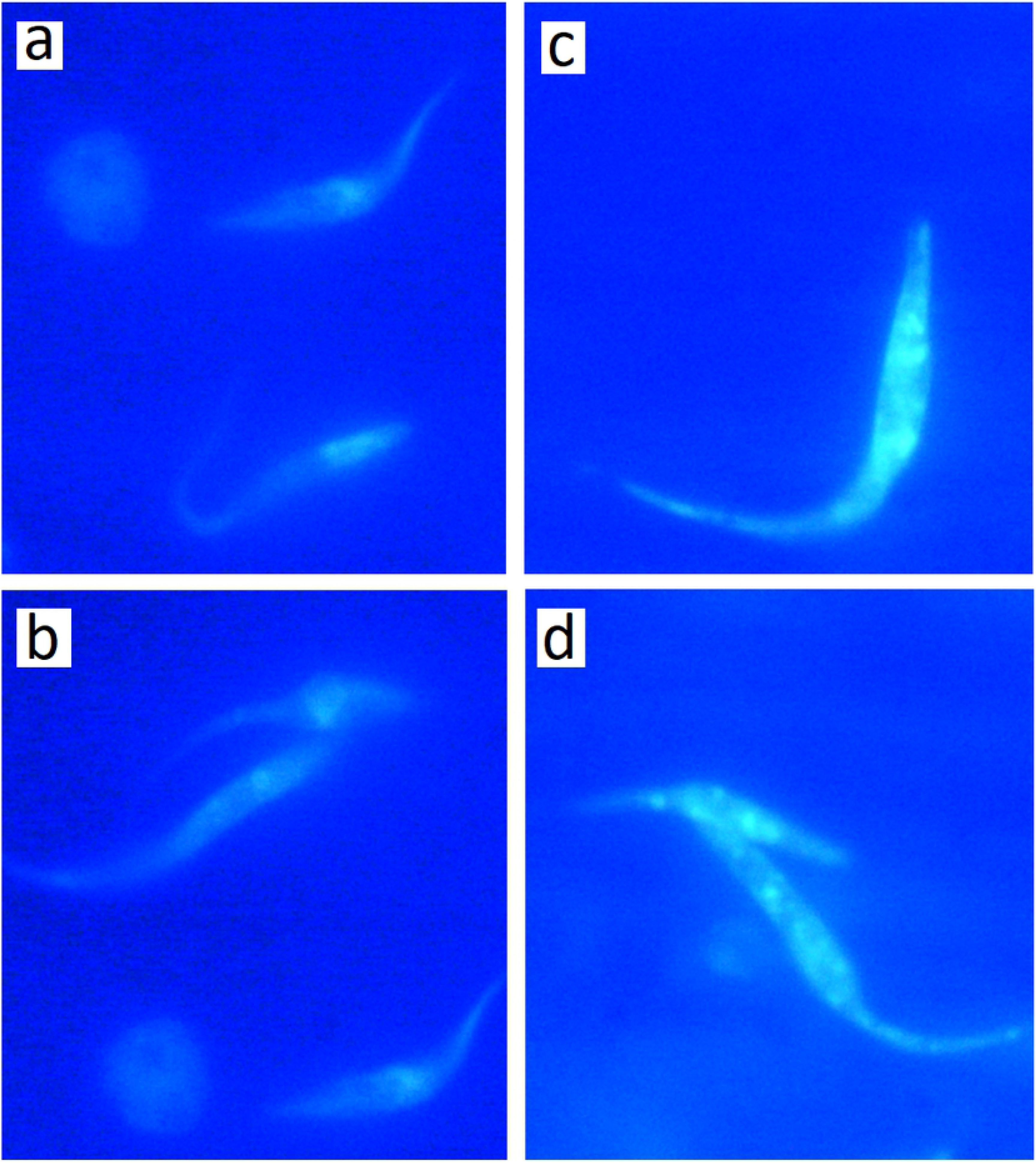
Autophagic induction by ST18 in *Trypanosoma cruzi* epimastigotes. Parasites were incubated with 0.05 mM MDC in PBS at 37 ° C for 10 minutes and observed in a fluorescent microscope Nikon Eclipse E 200 (100x). A and b: Control. c and d: *T. cruzi* epimastigotes treated 72 h with 40 μM ST18.Eclipse E 200 (100x). a and b: Control. c and d: *T. cruzi* epimastigotes treated 72 h with 40 μM ST18.

### Caspase-1

Infection with *T. cruzi* results in activation of caspase-1 and inflammasome formation. Inflammasome is indispensable for controlling parasitemia, host survival, and the onset of adaptive immune response [16]. In this sense, inflammasome activation is fully dependent on caspase-1. We evaluated the levels of active caspase-1 in U937 macrophages infected with *T. cruzi* after treatment with Nifurtimox, ST18 and TR4. In macrophages infected with trypanosomes and treated with ST18 spectrophotometric analysis showed a substantial increase in active caspase-1 compared to the control. In contrast no increase in caspase-1 was observed in samples of infected macrophages treated with Nifurtimox or TR4 and in uninfected macrophages treated with ST18. (Fig. 7).

**Fig 7.**
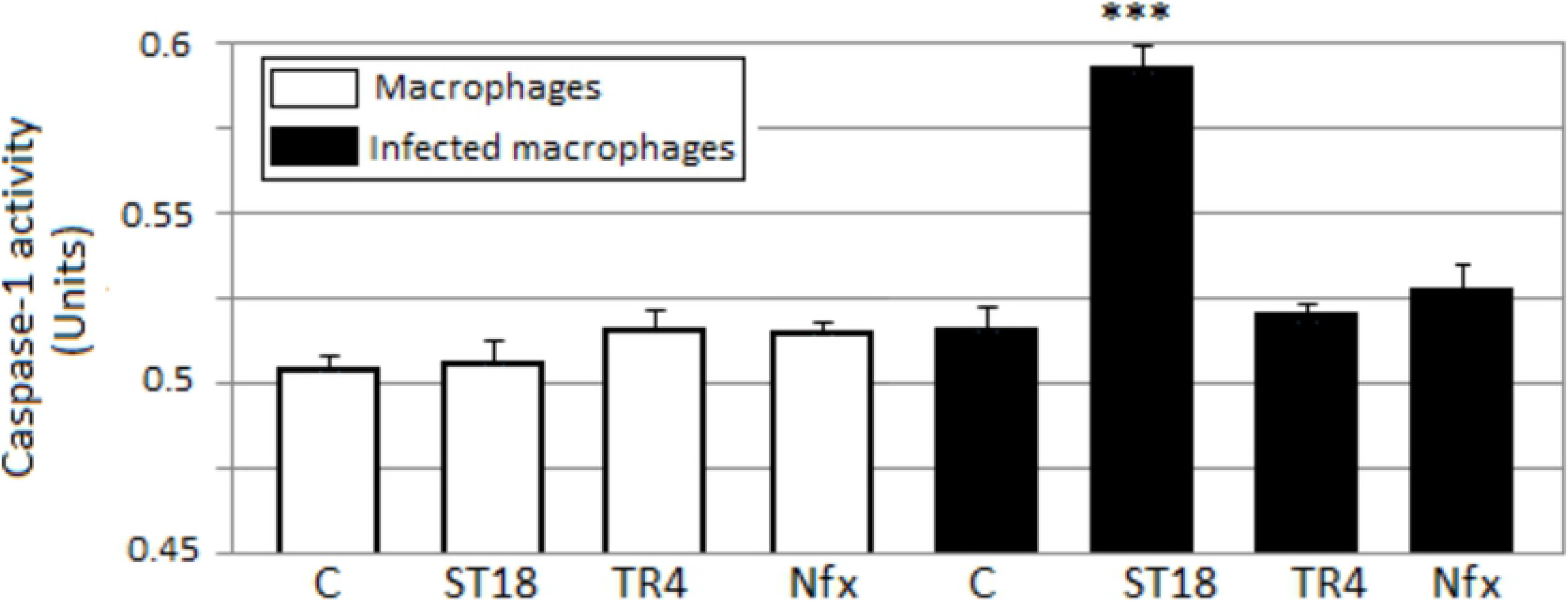
Caspase-1 activity. Levels of active caspase-1 in U937 macrophages infected with in *T. cruzi* epimastigotes after 48 hours of treatment with 50 μM of ST18, TR4 and Nifurtimox (Nfx). C = untreated control. Bars indicate the mean ± SE from four independent experiments. *p<0.05 vs control.

## Discussion

We evaluated the *in vitro* antiparasitic effects in *T. cruzi* epimastigotes of a series of *cis*- and *trans*-stilbenes bearing 3,5-dimetoxy motif at A phenyl ring and amino, methoxy and hydroxyl function at 2’, 3’- and/or 4’-position at B phenyl ring. Moreover, in an attempt to increase the chemical diversity of the compounds we studied a small series of terphenyl derivatives that notably do not bear the ethylene double bond that is the main reason for the chemical and metabolic instability of stilbenes [17,18]. Data were compared to those obtained with Nifurtimox which is the drug currently used for the treatment of Trypanosome infections. Among the stilbene series, ST18 bearing a 4’-methoxy function was the most active compound showing an IC50 = 4.6 μM ± 0.4. Regarding the therphenyl derivatives the best results were obtained with the trimethoxylated compound TR4 (IC50 = 30 ± 4.3) that is the *orto*-terphenyl analogue of ST18. Nifurtimox was more active than ST18 in *T. cruzi* epimastigotes but less active in intramacrophagic *T. cruzi* amastigotes.

The most interesting data observed in this study was the difference in the selectivity index value between ST18 and Nifurtimox. Nifurtimox is a drug with several adverse effects including mutagenic and tumorigenic effects [3]. ST18 has been described in the literature by different names, including resveratrol trimethyl ether (RTE) [19,20], MR-3 [21,22], M-5 [23], BTM-0521 [24], trimethoxy resveratrol [25], trimethylated resveratrol [26] and TMS [20,27]. It is a natural stilbene isolated from *Virola cuspidata* and *Virola elongata* bark [27,28]. Natural stilbenes have received increasing attention due to their potent antioxidant properties and their marked effects in the prevention of various oxidative stress associated diseases such as cancer [28]. A number of clinical trials using natural stilbenes such as resveratrol and pterostilbene have shown that they are therapeutically effective and pharmacologically safe because it showed no organ-specific or systemic toxicity [29–33]. Preclinical pharmacokinetic studies have shown that ST18 has appropriate pharmacokinetic profiles that make it a promising drug candidate for further pharmaceutical development [19]. It exhibited anti-proliferative and/or apoptosis-inductive activities in various cancer cells with a potency usually higher than resveratrol [20,23,34–36]. Moreover it has shown anti-inflammatory [37–40], gastro protective [41], and hepato-protective activities [26]. Here, we have demonstrated that ST18 showed a very low toxicity on normal and differentiated cells and the SI tested in *T. cruzi* parasites was higher than that calculated for Nifurtimox.

Several studies have shown that Nifurtimox induces production of reactive oxygen species (ROS) and subsequent apoptosis in neoplastic cells [42–44]. Although programmed cell death is very controversial in unicellular eukaryotes we observed that Nifurtimox caused a marked reduction in the average cell size of *T. cruzi* epimastigotes and a significant increase in the percentage of parasites positive to Annexin V. This compound did not cause in parasites an increase of MDC, an important marker of autophagy. In contrast ST18 produced a heterogeneous population characterized by parasites with low dimension and parasites with increased size and granularity. ST18 induced an increase of both Annexin V and MDC positive parasites.

Several works have reported the activation of autophagic process in Trypanosomatids during starvation responses and life cycle developments. Moreover, endoplasmic reticulum (ER) stress and anti-parasitic drugs can induce autophagy in *T. brucei* and *T. cruzi* [45–47]. In our experiments ST18 caused in *T. cruzi* both phosphatidylserine expression and dansylcadaverina staining suggesting that this compound could be capable to activate both apoptosis and autophagy.

Lim et al. [48] have obtained similar results in *T. brucei rhodesiense* using two piperidine alkaloids, (+)-spectaline and iso-6-spectaline. These compounds caused the formation of autophagic vacuoles were to monodansylcadaverine staining indicating the activation of the autophagic process. When trypanosomes were treated with piperidine alkaloids for 72 h they showed apoptotic aspects including phosphatidylserine exposure.

Several studies have demonstrated that autophagy and apoptosis communicate with each other to decides the fate of the cell during physiological and pathological conditions [49]. It has been supposed that, after the activation of stress or drug induced autophagy, when the stress condition increases towards a point of no return cells block autophagy and activate programmed cell death. Of interest, the analysis of cell cycle showed that both Nifurtimox and TR4 caused a decrease of parasites in G2M phase of cell cycle while ST18 determined an important block in G2M. A correlation between G2M block and autophagy activation has been observed in different experimental models but the precise mechanism by which microtubule targeting agents induce autophagic cell death is not known [50–53].

Finally, we observed that ST18, but not TR4 and Nifurtimox, induced a marked increase of active caspase-1 in *T. cruzi* infected macrophages. The capability of ST18 to activate caspase-1 in *T. cruzi* infected macrophages may, in part, explain the greater antiparasitic effect of ST18 than Nifurtimox in intramacrophagic trypanosomes. In fact, Yu et al. [54] demonstrated that canonical inflammasome activation triggers ROS production in macrophages in a caspase-1-dependent manner. Reactive oxygen species (ROS) protect the host against a large number of pathogenic microorganisms including trypanosome [55,56].

In conclusion, after testing 17 different compounds designed and synthesized previously by us, we selected a stilbene compound, ST18, endowed with a potent antiparasitic activity in *T. cruzi* epimastigotes and intracellular amastigotes. The antiparasitic activity of ST18 together to its ability to activate caspase-1 in infected macrophages and its low toxicity on normal cells makes this compound interesting for further biological and clinical studies in *T. cruzi*.

## Acknowledgments

This work is supported by grants from ‘Istituto Zooprofilattico Sperimentale della Sicilia”, Palermo, Italy code number: RC IZS SI 05/16.

## References

1. Shoemaker EA, Dale K, Cohn DA, Kelly MP, Zoerhoff KL, Batcho WE, et al. Gender and neglected tropical disease front-line workers: Data from 16 countries. PloS One. 2019;14: e0224925. doi:10.1371/journal.pone.0224925

2. Urbina, J. A., and R. Docampo. Specific chemotherapy of Chagas disease: controversies and advances. Trends Parasitol. 2003.

3. Castro JA, de Mecca MM, Bartel LC. Toxic side effects of drugs used to treat Chagas’ disease (American trypanosomiasis). Hum Exp Toxicol. 2006;25: 471–479. doi:10.1191/0960327106het653oa

4. Morais TR, Conserva GAA, Varela MT, Costa-Silva TA, Thevenard F, Ponci V, et al. Improving the drug-likeness of inspiring natural products - evaluation of the antiparasitic activity against Trypanosoma cruzi through semi-synthetic and simplified analogues of licarin A. Sci Rep. 2020;10: 5467. doi:10.1038/s41598-020-62352-w

5. Pizzirani D, Roberti M, Cavalli A, Grimaudo S, Di Cristina A, Pipitone RM, et al. Antiproliferative Agents That Interfere with the Cell Cycle at the G1→S Transition: Further Development and Characterization of a Small Library of Stilbene-Derived Compounds. ChemMedChem. 2008;3: 345–355. doi:10.1002/cmdc.200700258

6. Roberti M, Pizzirani D, Simoni D, Rondanin R, Baruchello R, Bonora C, et al. Synthesis and biological evaluation of resveratrol and analogues as apoptosis-inducing agents. J Med Chem. 2003;46: 3546–3554. doi:10.1021/jm030785u

7. Roberti M. Identification of a Terphenyl Derivative that Blocks the Cell Cycle in the G0−G1 Phase and Induces Differentiation in Leukemia Cells | Journal of Medicinal Chemistry. 2006 [cited 2 Feb 2021]. Available: https://pubs.acs.org/doi/pdf/10.1021/jm060253o

8. Tolomeo M, Roberti M, Scapozza L, Tarantelli C, Giacomini E, Titone L, et al. TTAS a new stilbene derivative that induces apoptosis in Leishmania infantum. Exp Parasitol. 2013;133: 37–43. doi:10.1016/j.exppara.2012.10.006

9. Castelli G, Bruno F, Vitale F, Roberti M, Colomba C, Giacomini E, et al. In vitro antileishmanial activity of trans-stilbene and terphenyl compounds. Exp Parasitol. 2016;166: 1–9. doi:10.1016/j.exppara.2016.03.007

10. Bruno F, Castelli G, Vitale F, Giacomini E, Roberti M, Colomba C, et al. Effects of trans-stilbene and terphenyl compounds on different strains of Leishmania and on cytokines production from infected macrophages. Exp Parasitol. 2018;184: 31–38. doi:10.1016/j.exppara.2017.11.004

11. Castelli G, Galante A, Lo Verde V, Migliazzo A, Reale S, Lupo T, et al. Evaluation of two modified culture media for Leishmania infantum cultivation versus different culture media. J Parasitol. 2014;100: 228–230. doi:10.1645/13-253.1

12. Kim S, Ko H, Park JE, Jung S, Lee SK, Chun Y-J. Design, synthesis, and discovery of novel trans-stilbene analogues as potent and selective human cytochrome P450 1B1 inhibitors. J Med Chem. 2002;45: 160–164. doi:10.1021/jm010298j

13. Munafó DB, Colombo MI. A novel assay to study autophagy: regulation of autophagosome vacuole size by amino acid deprivation. J Cell Sci. 2001;114: 3619–3629.

14. V J, R P, Ma S, N G. Natural programmed cell death in T. cruzi epimastigotes maintained in axenic cultures. In: Journal of cellular biochemistry [Internet]. 15 Oct 2008 [cited 15 Feb 2021]. doi:10.1002/jcb.21864

15. Biederbick A, Kern HF, Elsässer HP. Monodansylcadaverine (MDC) is a specific in vivo marker for autophagic vacuoles. Eur J Cell Biol. 1995;66: 3–14.

16. paroli. Frontiers | NLRP3 Inflammasome and Caspase-1/11 Pathway Orchestrate Different Outcomes in the Host Protection Against Trypanosoma cruzi Acute Infection | Immunology. [cited 2 Feb 2021]. Available: https://www.frontiersin.org/articles/10.3389/fimmu.2018.00913/full

17. Metzler, M. Metabolic activation of diethylstilbestrol: Indirect evidence for the formation of a stilbene oxide intermediate in hamster and rat. Biochem Pharmacol. 1975;24: 1449–1453. doi:10.1016/0006-2952(75)90373-1

18. Neumann HG, Metzler M, Töpner W. Metabolic activation of diethylstilbestrol and aminostilbene-derivatives. Arch Toxicol. 1977;39: 21–30. doi:10.1007/BF00343272

19. Hs L, Pc H. Preclinical pharmacokinetic evaluation of resveratrol trimethyl ether in sprague-dawley rats: the impacts of aqueous solubility, dose escalation, food and repeated dosing on oral bioavailability. In: Journal of pharmaceutical sciences [Internet]. Oct 2011 [cited 15 Feb 2021]. doi:10.1002/jps.22588

20. Tt W, Nw S, Ys K, Cs M, Am R. Differential effects of resveratrol and its naturally occurring methylether analogs on cell cycle and apoptosis in human androgen-responsive LNCaP cancer cells. In: Molecular nutrition & food research [Internet]. Mar 2010 [cited 15 Feb 2021]. doi:10.1002/mnfr.200900143

21. Cj W, Yt Y, Ct H, Gc Y. Mechanisms of apoptotic effects induced by resveratrol, dibenzoylmethane, and their analogues on human lung carcinoma cells. In: Journal of agricultural and food chemistry [Internet]. 24 Jun 2009 [cited 15 Feb 2021]. doi:10.1021/jf900531m

22. Yt Y, Cj W, Ct H, Gc Y. Resveratrol analog-3,5,4’-trimethoxy-trans-stilbene inhibits invasion of human lung adenocarcinoma cells by suppressing the MAPK pathway and decreasing matrix metalloproteinase-2 expression. In: Molecular nutrition & food research [Internet]. Mar 2009 [cited 15 Feb 2021]. doi:10.1002/mnfr.200800123

23. Bader Y, Madlener S, Strasser S, Maier S, Saiko P, Stark N, et al. Stilbene analogues affect cell cycle progression and apoptosis independently of each other in an MCF-7 array of clones with distinct genetic and chemoresistant backgrounds. Oncol Rep. 2008;19: 801–810. doi:10.3892/or.19.3.801

24. Lu M, Liu B, Xiong H, Wu F, Hu C, Liu P. Trans-3,5,4’-trimethoxystilbene reduced gefitinib resistance in NSCLCs via suppressing MAPK/Akt/Bcl-2 pathway by upregulation of miR-345 and miR-498. J Cell Mol Med. 2019;23: 2431–2441. doi:10.1111/jcmm.14086

25. Sj D, K L, Am R, S D, Cs M, Ad P, et al. Trimethoxy-resveratrol and piceatannol administered orally suppress and inhibit tumor formation and growth in prostate cancer xenografts. In: The Prostate [Internet]. Aug 2013 [cited 15 Feb 2021]. doi:10.1002/pros.22657

26. H R, M S, V T, V P-A, P M. Resveratrol and trimethylated resveratrol protect from acute liver damage induced by CCl4 in the rat. J Appl Toxicol JAT. 2008;28: 147–155. doi:10.1002/jat.1260

27. B L, Xj L, Zb Y, Jj Z, Tb L, Xj Z, et al. Inhibition of NOX/VPO1 pathway and inflammatory reaction by trimethoxystilbene in prevention of cardiovascular remodeling in hypoxia-induced pulmonary hypertensive rats. In: Journal of cardiovascular pharmacology [Internet]. Jun 2014 [cited 15 Feb 2021]. doi:10.1097/FJC.0000000000000082

28. Sirerol JA, Rodríguez ML, Mena S, Asensi MA, Estrela JM, Ortega AL. Role of Natural Stilbenes in the Prevention of Cancer. Oxid Med Cell Longev. 2016;2016: 3128951. doi:10.1155/2016/3128951

29. Almeida L, Vaz-da-Silva M, Falcão A, Soares E, Costa R, Loureiro AI, et al. Pharmacokinetic and safety profile of trans-resveratrol in a rising multiple-dose study in healthy volunteers. Mol Nutr Food Res. 2009;53 Suppl 1: S7–15. doi:10.1002/mnfr.200800177

30. la Porte C, Voduc N, Zhang G, Seguin I, Tardiff D, Singhal N, et al. Steady-State pharmacokinetics and tolerability of trans-resveratrol 2000 mg twice daily with food, quercetin and alcohol (ethanol) in healthy human subjects. Clin Pharmacokinet. 2010;49: 449–454. doi:10.2165/11531820-000000000-00000

31. Boocock DJ, Faust GES, Patel KR, Schinas AM, Brown VA, Ducharme MP, et al. Phase I Dose Escalation Pharmacokinetic Study in Healthy Volunteers of Resveratrol, a Potential Cancer Chemopreventive Agent. Cancer Epidemiol Prev Biomark. 2007;16: 1246–1252. doi:10.1158/1055-9965.EPI-07-0022

32. Brown VA, Patel KR, Viskaduraki M, Crowell JA, Perloff M, Booth TD, et al. Repeat dose study of the cancer chemopreventive agent resveratrol in healthy volunteers: safety, pharmacokinetics, and effect on the insulin-like growth factor axis. Cancer Res. 2010;70: 9003–9011. doi:10.1158/0008-5472.CAN-10-2364

33. Patel KR, Brown VA, Jones DJL, Britton RG, Hemingway D, Miller AS, et al. Clinical pharmacology of resveratrol and its metabolites in colorectal cancer patients. Cancer Res. 2010;70: 7392–7399. doi:10.1158/0008-5472.CAN-10-2027

34. D S, M R, Fp I, E A, S A, P M, et al. Stilbene-based anticancer agents: resveratrol analogues active toward HL60 leukemic cells with a non-specific phase mechanism. In: Bioorganic & medicinal chemistry letters [Internet]. 15 Jun 2006 [cited 15 Feb 2021]. doi:10.1016/j.bmcl.2006.03.028

35. V C, R C, L L, S S, C S, C T. Antiproliferative activity of methylated analogues of E- and Z-resveratrol. In: Zeitschrift fur Naturforschung. C, Journal of biosciences [Internet]. Apr 2007 [cited 15 Feb 2021]. doi:10.1515/znc-2007-3-406

36. Pan M-H, Gao J-H, Lai C-S, Wang Y-J, Chen W-M, Lo C-Y, et al. Antitumor activity of 3,5,4′-trimethoxystilbene in COLO 205 cells and xenografts in SCID mice. Mol Carcinog. 2008;47: 184–196. doi:10.1002/mc.20352

37. Yuan Q, Peng J, Liu S-Y, Wang C-J, Xiang D-X, Xiong X-M, et al. Inhibitory effect of resveratrol derivative BTM-0512 on high glucose-induced cell senescence involves dimethylaminohydrolase/asymmetric dimethylarginine pathway. Clin Exp Pharmacol Physiol. 2010;37: 630–635. doi:10.1111/j.1440-1681.2010.05368.x

38. Yh D, D A, Hq H, N W, N Y, Yt W, et al. Inhibition of TNF-α-mediated endothelial cell-monocyte cell adhesion and adhesion molecules expression by the resveratrol derivative, trans-3,5,4’-trimethoxystilbene. In: Phytotherapy research: PTR [Internet]. Mar 2011 [cited 15 Feb 2021]. doi:10.1002/ptr.3279

39. Xl M, Jy Y, Gl C, Lh W, Lj Z, S W, et al. Effects of resveratrol and its derivatives on lipopolysaccharide-induced microglial activation and their structure-activity relationships. In: Chemico-biological interactions [Internet]. 7 Oct 2008 [cited 15 Feb 2021]. doi:10.1016/j.cbi.2008.04.015

40. Effects of resveratrol-related hydroxystilbenes on the nitric oxide production in macrophage cells: structural requirements and mechanism of action. Life Sci. 2002;71: 2071–2082. doi:10.1016/S0024-3205(02)01971-9

41. Farzaei MH, Abdollahi M, Rahimi R. Role of dietary polyphenols in the management of peptic ulcer. World J Gastroenterol. 2015;21: 6499–6517. doi:10.3748/wjg.v21.i21.6499

42. Sholler GLS, Brard L, Straub JA, Dorf L, Illyene S, Koto K, et al. Nifurtimox Induces Apoptosis of Neuroblastoma Cells in vitro and in vivo. J Pediatr Hematol Oncol. 2009;31: 187–193. doi:10.1097/MPH.0b013e3181984d91

43. Du M, Zhang L, Scorsone KA, Woodfield SE, Zage PE. Nifurtimox Is Effective Against Neural Tumor Cells and Is Synergistic with Buthionine Sulfoximine. Sci Rep. 2016;6: 27458. doi:10.1038/srep27458

44. Koto KS, Lescault P, Brard L, Kim K, Singh RK, Bond J, et al. Antitumor activity of nifurtimox is enhanced with tetrathiomolybdate in medulloblastoma. Int J Oncol. 2011;38: 1329–1341. doi:10.3892/ijo.2011.971

45. Goldshmidt H, Matas D, Kabi A, Carmi S, Hope R, Michaeli S. Persistent ER stress induces the spliced leader RNA silencing pathway (SLS), leading to programmed cell death in Trypanosoma brucei. PLoS Pathog. 2010;6: e1000731. doi:10.1371/journal.ppat.1000731

46. Uzcátegui NL, Carmona-Gutiérrez D, Denninger V, Schoenfeld C, Lang F, Figarella K, et al. Antiproliferative effect of dihydroxyacetone on Trypanosoma brucei bloodstream forms: cell cycle progression, subcellular alterations, and cell death. Antimicrob Agents Chemother. 2007;51: 3960–3968. doi:10.1128/AAC.00423-07

47. Delgado M, Anderson P, Garcia-Salcedo JA, Caro M, Gonzalez-Rey E. Neuropeptides kill African trypanosomes by targeting intracellular compartments and inducing autophagic-like cell death. Cell Death Differ. 2009;16: 406–416. doi:10.1038/cdd.2008.161

48. Lim KT, Yeoh CY, Zainuddin Z, Ilham Adenan M. (+)-Spectaline and Iso-6-Spectaline Induce a Possible Cross-Talk between Autophagy and Apoptosis in Trypanosoma brucei rhodesiense. Trop Med Infect Dis. 2019;4. doi:10.3390/tropicalmed4030098

49. Ojha R, Ishaq M, Singh SK. Caspase-mediated crosstalk between autophagy and apoptosis: Mutual adjustment or matter of dominance. J Cancer Res Ther. 2015;11: 514–524. doi:10.4103/0973-1482.163695

50. Acharya BR, Bhattacharyya S, Choudhury D, Chakrabarti G. The microtubule depolymerizing agent naphthazarin induces both apoptosis and autophagy in A549 lung cancer cells. Apoptosis Int J Program Cell Death. 2011;16: 924–939. doi:10.1007/s10495-011-0613-1

51. Reunanen H, Marttinen M, Hirsimäki P. Effects of griseofulvin and nocodazole on the accumulation of autophagic vacuoles in Ehrlich ascites tumor cells. Exp Mol Pathol. 1988;48: 97–102. doi:10.1016/0014-4800(88)90048-2

52. Kuo P-L, Hsu Y-L, Cho C-Y. Plumbagin induces G2-M arrest and autophagy by inhibiting the AKT/mammalian target of rapamycin pathway in breast cancer cells. Mol Cancer Ther. 2006;5: 3209–3221. doi:10.1158/1535-7163.MCT-06-0478

53. Kondo Y, Kanzawa T, Sawaya R, Kondo S. The role of autophagy in cancer development and response to therapy. Nat Rev Cancer. 2005;5: 726–734. doi:10.1038/nrc1692

54. Yu J, Nagasu H, Murakami T, Hoang H, Broderick L, Hoffman HM, et al. Inflammasome activation leads to Caspase-1-dependent mitochondrial damage and block of mitophagy. Proc Natl Acad Sci U S A. 2014;111: 15514–15519. doi:10.1073/pnas.1414859111

55. Richard M. Locksley Seymour J. Klebanoff. Oxygen-dependent microbicidal systems of phagocytes and host defense against intracellular protozoa - Locksley - 1983 - Journal of Cellular Biochemistry - Wiley Online Library. [cited 2 Feb 2021]. Available: https://onlinelibrary.wiley.com/doi/abs/10.1002/jcb.240220306

56. Nathan C, Nogueira N, Juangbhanich C, Ellis J, Cohn Z. Activation of macrophages in vivo and in vitro. Correlation between hydrogen peroxide release and killing of Trypanosoma cruzi. J Exp Med. 1979;149: 1056–1068. doi:10.1084/jem.149.5.1056

